# METTL13 Promotes Pre-Leukemic Transformation and the Development of Pediatric Leukemia

**DOI:** 10.1101/2025.08.29.673071

**Authors:** Sabina Enlund, Chae-Eun Lim, Isabella Hoang, Sonali Joshi, Amanda Ramilo Amor, Cecilia Thomsson, Indranil Sinha, Shahrzad Shirazi Fard, Anna Nilsson, Ola Hermanson, Qingfei Jiang, Frida Holm

## Abstract

Post-transcriptional RNA modifications, such as N6-methyladenosine (m6A) methylation and adenosine to inosine (A-to-I) editing, are critical regulators of hematopoietic stem cell (HSC) self-renewal and differentiation, yet their precise contributions to malignant transformation are not fully elucidated. In this study, we uncovered the epitranscriptomic landscape caused by knockdown of genes from the methyltransferase (METTL)-family in hematopoietic stem and progenitor cells (HSPCs). We identified both converging and distinct roles of METTL3 and METTL14, known members of the m6A writer complex, as well as orphan gene METTL13. Notably, METTL13 was uniquely upregulated by adenosine deaminase acting on RNA 1 (ADAR1) overexpression, while other METTL genes were downregulated. Knockdown of METTL13 altered the expression of multiple genes involved in oncogenic development in HSPCs. Furthermore, METTL13 was associated with a high-risk profile in pediatric T-cell acute lymphoblastic leukemia (T-ALL), and functional studies confirmed that METTL13 is required for T-ALL cell proliferation and survival both *in vitro* and *in vivo*. Collectively, our results indicate a previously unrecognized, oncogenic role for METTL13 in pre-leukemic transformation and T-ALL pathogenesis.

**Significance:** In this study we uncovered a novel regulatory link between ADAR1 and the METTL-family of RNA methyltransferases in hematopoietic stem cells. Overexpression of ADAR1 uniquely upregulated METTL13 while suppressing other METTL genes. Loss of orphan gene METTL13 affected proliferation, apoptosis and p53 signaling in hematopoietic stem cells. Furthermore, loss of METTL13 suppressed cell proliferation and survival in pediatric T-cell acute lymphoblastic leukemia. Our findings suggest a potential role for METTL13 in pre-leukemia transformation and oncogenic development.

## Introduction

Epitranscriptomic alterations, which comprise numerous post-transcriptional RNA modifications, have been subjected to extensive research for their implications in disease development and cancer progression over the last decade (1). RNA modifications play a crucial role in the cellular response to environmental cues, regulating cell survival, differentiation and migration. Aberrant activity of these RNA modifications can contribute to malignant transformation of normal cells and has been linked to a wide variety of cancers (2).

Among RNA modifications, N6-methyladenosine (m6A) methylation and adenosine to inosine (A-to-I) editing are the most abundant, both targeting adenosines on mRNA to alter its fate and function (3). A-to-I editing is achieved by adenosine deaminase acting on RNA 1 (ADAR1) and has been widely studied for its role in regulating innate immunity and maintaining normal hematopoietic stem cells (HSCs) (4–6). Aberrant A-to-I RNA editing leads to many consequences, including impaired stem cell fitness, premature aging, and cancer development (7–9). Malignant overexpression of ADAR1 has been reported in over 20 types of cancers, including chronic myeloid leukemia and pediatric T-cell acute lymphoblastic leukemia (T-ALL) (7, 10).

RNA methylation by the m6A complex is a dynamic and reversible process mediated by a large network of proteins, many of which are not completely characterized (3, 11–13). Methylation is accomplished by a group of methyltransferases (METTLs), predominantly METTL3 and METTL14, which make up the core of the writer complex (14). The METTL3-METTL14 complex further interacts with proteins like WTAP and METTL16 for specific recruitment of RNA. The reversible capacity of m6A modifications is achieved by two known erasers, fat mass and obesity-associated (FTO) and demethylase AlkB homolog 5 (ALKBH5) (13, 15–17). Different readers, such as YTH-containing proteins and heterogeneous nuclear ribonucleoproteins (HNRNPs), recognize and interpret m6A sites on mRNA transcripts, ultimately determining the transcript fate and downstream effects (13, 18–20). Modifications by the m6A complex have diverse outcomes, affecting many aspects of mRNA metabolism, including mRNA stability, splicing and translation (21–24).

Abnormal activity of m6A methylation has been associated with many diseases, including neurological and autoimmune disorders, as well as impaired stem cell differentiation (2, 16, 25–30). Interestingly, many of m6A functions overlap with those of ADAR1. Both highly regulated m6A methylation and A-to-I editing are required for HSC self-renewal and differentiation (4–6, 8, 31–36). Similar to malignant A-to-I editing, alterations to m6A modifications promote the development of both solid tumors and hematological cancers, including acute myeloid leukemia (37), (36, 38–40). Recent studies have also concluded a critical role for m6A methylation in maintaining stem cell-like capacities of leukemia stem cells (LSCs), responsible for both disease initiation and relapse. Aberrant activity of m6A complex was shown to promote LSC maintenance, thus potentiating therapy resistance and tumor metastasis (41).

These two types of RNA modifications are known to regulate HSC fitness separately, but little is known about how these pathways interact to fine-tune cellular functions in normal HSCs and in the transformation to LSCs. Studies have shown that m6A writer METTL3 and reader YTHDF1 can directly introduce m6A in ADAR1 mRNA, thereby promoting ADAR1 expression and A-to-I RNA editing (42, 43). Both m6A and A-to-I RNA modifications target adenosine bases, although it is still debated whether they compete for the exact same adenosine. A-to-I RNA editing occurs mainly in inverted Alu sequences enriched in intronic and 3’UTR regions, while m6A are located within highly conserved RRACH sequences within 3’UTR and exons (34, 44). Whether these two RNA-modifying processes functionally interact to promote malignant transformation and leukemia development is yet to be defined.

In this study, we aimed to unravel the role of these RNA modifications in HSC maintenance and in the development of hematological malignancies. We hypothesized that m6A methylation and A-to-I editing, two essential cellular machineries, crosstalk and coordinate their activities, leading to malignant transformation of HSCs. We report that enforced ADAR1 overexpression in healthy hematopoietic stem and progenitor cells (HSPC) led to differential expression of the METTL-family and other m6A-related components. Most m6A-related genes were downregulated following ADAR1 overexpression, except for METTL13, an orphan METTL-family member whose function in HSCs and LSCs has not yet been reported. Knockdown of METTL3, METTL13 and METTL14 affected numerous overlapping genes involved in inflammatory signaling pathways and cell survival in HSPCs. Furthermore, functional enrichment analysis following METTL13 knockdown revealed dysregulation of pathways involved in cancer progression as well as disease ontologies related to hematological cancer. Finally, we investigated the role of METTL13 in pediatric acute lymphoblastic leukemia (ALL). Our results revealed a potential association between increased METTL13 expression and a worse prognosis in both T-ALL and B-cell ALL (B-ALL) patients. Loss of METTL13 led to arrested cell proliferation and reduced cell viability in T-ALL cells, both *in vivo* and *in vitro*. RNA-sequencing analysis of METTL13 knockdown in T-ALL cells revealed a loss of many signaling pathways, several that are central for controlling proliferation and DNA replication. Taken together, our findings reveal consequences of alterations to genes in the METTL-family in the context of disease development and recognizes an emerging role of METTL13 in hematological malignancies.

## Results

### ADAR1 overexpression altered expression of the METTL-family and other components of the m6A complex

To investigate how ADAR1 regulates the m6A complex, we examined effects following lentiviral overexpression of ADAR1 in human CD34^+^ HSPCs (available from previous studies, at BioProject: PRJNA319866) (Fig. S1A). A total of 6,865 genes were significantly differentially expressed in ADAR1 overexpressed HSPCs compared to the backbone control (pCDH), with the majority (> 85%) of genes being downregulated (Fig. 1A, B, Supplemental table 1). By focusing on the m6A regulatory gene network, we found that ADAR1 overexpression led to downregulation of the main m6A writer complex (METTL3, METTL14 and WTAP) (28, 45, 46), m6A erasers (FTO and ALKBH5), and readers (HNRNPC and YTHDF1) (Fig. 1D, 1E). Other suppressed methyltransferase-like genes included METTL7A, a thiol methyltransferase(47); METTL9, a methyltransferase for 1-methylhistidine (48); and METTL16, which functions as both a non-conical nuclear m6A writer and cytosolic translation-initiator (49) (Figure SF1B). The only significantly upregulated gene from the METTL-family was METTL13. Together, these results indicate potential negative regulation of ADAR1 on several members of the METTL-family, at least on a transcriptional level, distinguishing METTL13 as the only METTL gene upregulated by ADAR1. Since METTL3 and METTL14 comprise the main m6A writer complex, this suggests ADAR1 may suppress m6A modifications.

**Figure 1.**
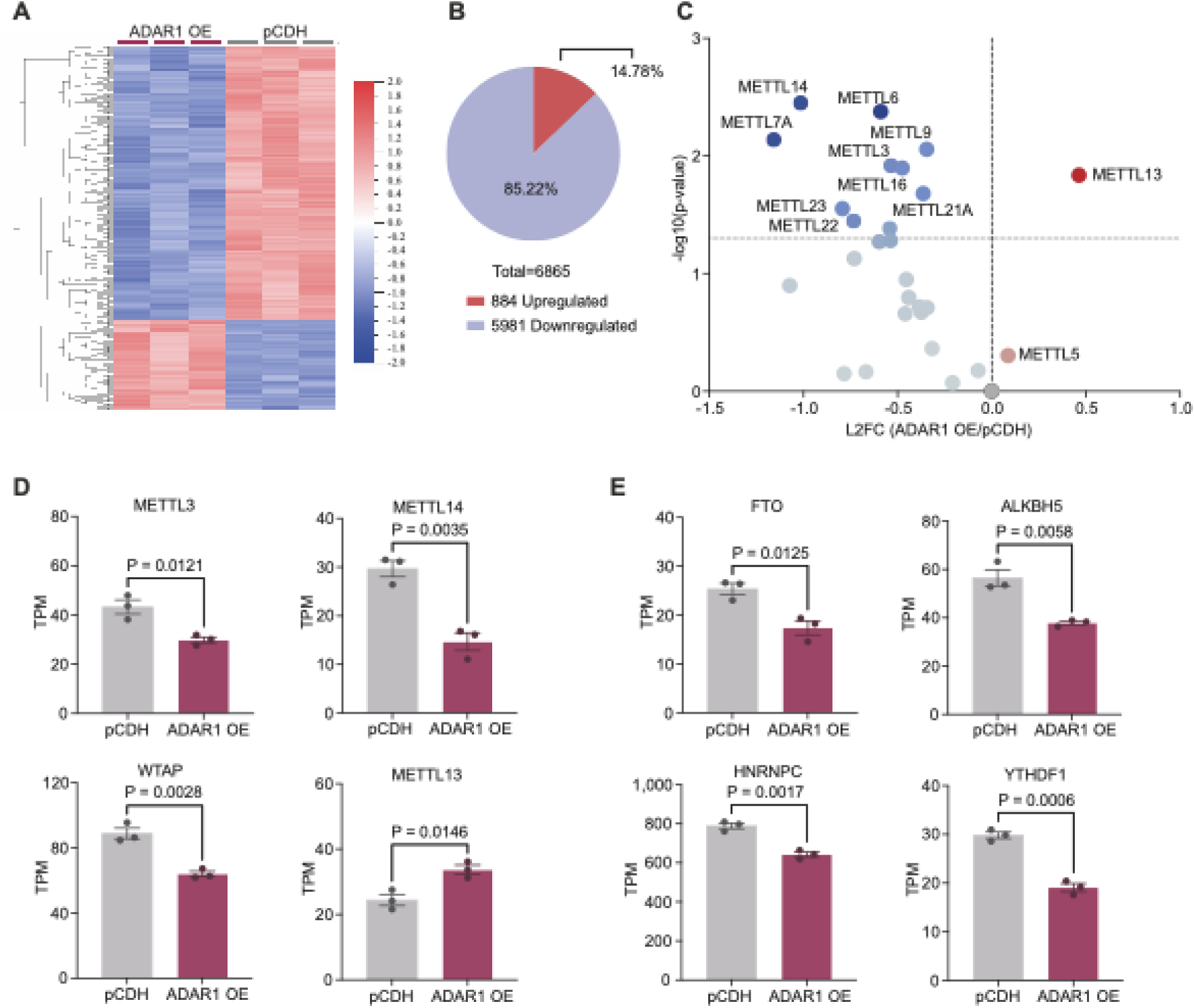
ADAR1 overexpression altered expression levels of METTL genes and other members of the m6A complex. A) Heat map of the top 500 differentially expressed genes in ADAR1 overexpressed human CD34^+^ HSPCs (ADAR1 OE, n=3) compared to the backbone control (pCDH, n=3). Created in Qlucore Omics Explorer, significance was calculated using unpaired two-tailed t-test, p<0.05. B) Distribution of differentially expressed genes in ADAR1 overexpressed HSPCs compared to the pCDH control. Significance was calculated using unpaired two-tailed t-test, p<0.05. C) Volcano plot of dysregulated genes from the METTL-family following ADAR1 overexpression. Significance was calculated usin unpaired two-tailed t-test, results are displayed as L2FC (ADAR OE/pCDH) and negative log10 p-value. D) Expression of genes from the m6A writer complex (METTL3, METTL14 and WTAP) and from the METTL-family (METTL13) in ADAR1 overexpressed cells compared to compared to the pCDH control. Significance was calculated using unpaired two-tailed t-test, results are displayed as TPM, mean ± SEM. E) Significantly differentially expressed m6A erasers FTO and ALKBH5 and readers HNRNPC and YTHDF1 in ADAR1 overexpressed cells compared to compared to the pCDH control. Significance was calculated using unpaired two-tailed t-test, results are displayed as TPM, mean ± SEM.

### Overlapping and distinct gene regulation programs by METTL3, METTL9, METTL13 and METTL14 in HSPCs

We systematically studied the gene expression landscape of several m6A family members affected by ADAR1 in human CD34^+^ HSPCs. Although the role of METTL3 and METTL14 are well established in HSC maintenance, METTL5, 9, and 13 have not been extensively studied in this cellular context. To achieve this, lentiviral knockdown of METTL3, METTL5, METTL9, METTL13 and METTL14 was performed in CD34+ HSPCs (Fig. SF2A, SF2B). Exploration using principal component analysis (PCA) revealed three potential outlier samples, which were excluded in the subsequent analysis (Fig. SF2C). Knockdown of METTL3 and METTL14, the core m6A writer complex, clustered together and were distinct from the control (Fig. 2A). Both shMETTL5 and shMETTL9 grouped together and showed similar effects on gene expression as the control, suggesting that METTL5 and METTL9 may have less effect on a transcriptional level, although knockdown efficacy requires improvement for further conclusions (Fig. 2A, 2B). Knockdown of METTL13 appeared to have similar effects on gene expression patterns as METTL3 and METTL14, yet shMETTL13 HSPCs formed their own cluster away from other METTLs and the control in PCA (Fig. 2A).

**Figure 2.**
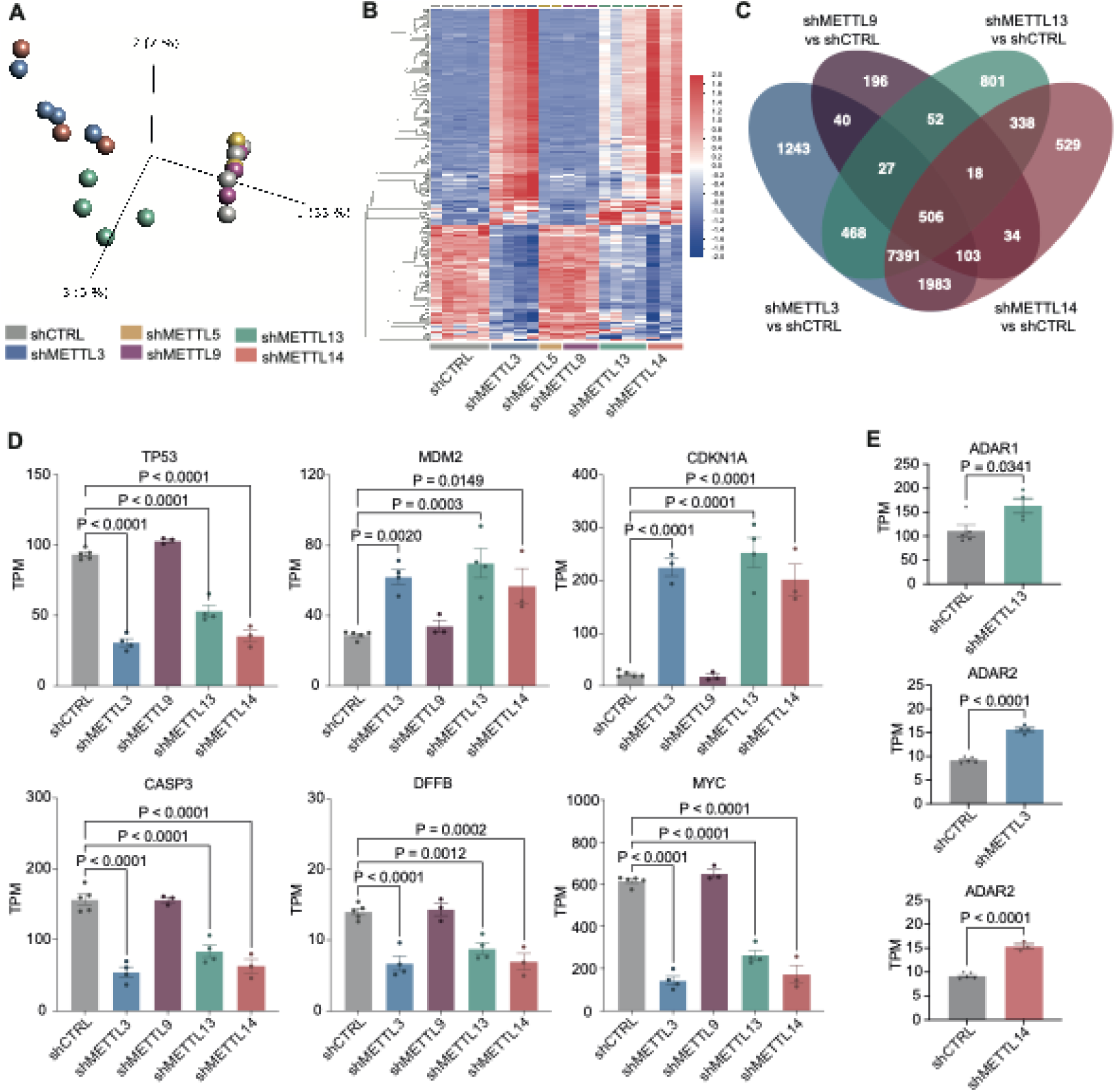
Overlapping and distinct gene regulation by METTL3, METTL9, METTL13 and METTL14 in HSPCs. A) PCA plot of METTL3 (shMETTL3, n=4), METTL5 (shMETTL5, n=2), METTL9 (shMETTL9, n=3), METTL13 (shMETTL13, n=4) and METTL14 (shMETTL14, n=3) knockdown in human CD34^+^ HSPCs compared to the backbone control (shCTRL, n=5). Created in Qlucore Omics Explorer. B) Heatmap of shMETTL3, shMETTL5, shMETTL9, shMETTL13 and shMETTL14 in HSPCs compared to shCTRL. Created in Qlucore Omics Explorer, significance was calculated using multi-group ANOVA, q<0.1, SD<0.05. C) Venn diagram of differentially expressed genes in shMETTL3, shMETTL9, shMETTL13 and shMETTL14 in HSPCs compared to shCTRL. Significance was calculated using unpaired two-tailed t-test, p<0.05, in each shMETTL compared to shCTRL. D) Dysregulated genes (TP53, MDM2, CDKN1A, CASP3, DFFB and c-MYC) in shMETTL3, shMETTL9, shMETTL13 and shMETTL14 compared to shCTRL. Significance was calculated using ordinary one-way ANOVA with multiple comparisons compared to shCTRL, as well as Dunnett correction, results are displayed as TPM, mean ± SEM. E) Differentially expressed genes from the ADAR family in shMETTL3, shMETTL13 and shMETTL14 compared to shCTRL. Significance was calculated using unpaired two-tailed t-test, results are displayed as TPM, mean ± SEM.

Next, we analyzed differentially expressed genes in each individual METTL knockdown condition compared to the control (Fig. 2C, Supplemental table 2). We excluded METTL5 due to low power following RNA quality control. Knockdown of METTL3, METTL13 and METTL14 shared many common differentially expressed genes (a total of 7391 genes), while knockdown of METTL3 and METTL14 alone shared only 1,983 genes. Knockdown of METTL9 affected fewer than 1,000 genes, while knockdown of METTL3, METTL13 and METTL14 altered expression of close to 10,000 genes independently, with most genes being downregulated (Fig. SF2D).

Our results indicate that the gene expression program of METTL13 substantially overlaps with that of m6A writer complex. To investigate this further, we carefully examined the overlapping target genes in HSPCs with depleted METTL3, METTL13 and METTL14. Reduced expression of tumor suppressor TP53 and upregulation of its negative regulator MDM2 were reported (Fig. 2D) (Fig. 2D) (50). Moreover, CDKN1A (codes for p21), an effector protein downstream of p53, essential for controlling cell cycle arrest, was upregulated (Fig. SD) (51). Genes involved in apoptotic signaling were inhibited, including CASP3 and DFFB, which are involved in degrading DNA during apoptosis (52). Additionally, expression of oncogene c-MYC was suppressed by METTL knockdown (53). Since m6A modification was reported to promote ADAR1 activity (42), we examined if loss of METTL would alter ADAR1 expression. We found that only knockdown of METTL13 led to increased ADAR1 expression (Fig. 2E). METTL3 and METTL14 knockdown caused an upregulation of ADAR2, which is mainly expressed and active within brain tissue, rather than the hematopoietic system (7). Taken together, these data suggest that METTL13 may converge with METTL3 and METTL14 in regulating gene expression in human HSPCs. In contrast, the separate PCA clustering and unique differentially expressed genes indicates that METTL13 may have some unique properties separate from METTL3 and METTL14, yet to be uncovered.

### METTL3, METTL13 and METTL14 converged in regulating immune signaling while METTL13 caused distinct effects on apoptosis and p53 regulation

To uncover both the shared and distinct cellular processes and signaling pathways controlled by METTL3, METTL9, METTL13 and METTL14 in HSPCs, we performed gene set enrichment analysis (GSEA) on each shMETTL compared to shCTRL, utilizing the WIKI Pathway, Reactome and KEGG libraries. Enrichment analysis revealed the most unique altered pathways following METTL13 knockdown across all three libraries (Fig. 3A, SF3A, SF3B, SF3C). Most pathways that were altered by the m6A writer complex (METTL3 and METTL14) were also altered by METTL13, while METTL3 and METTL14 knockdown alone caused minimal overlap.

**Figure 3.**
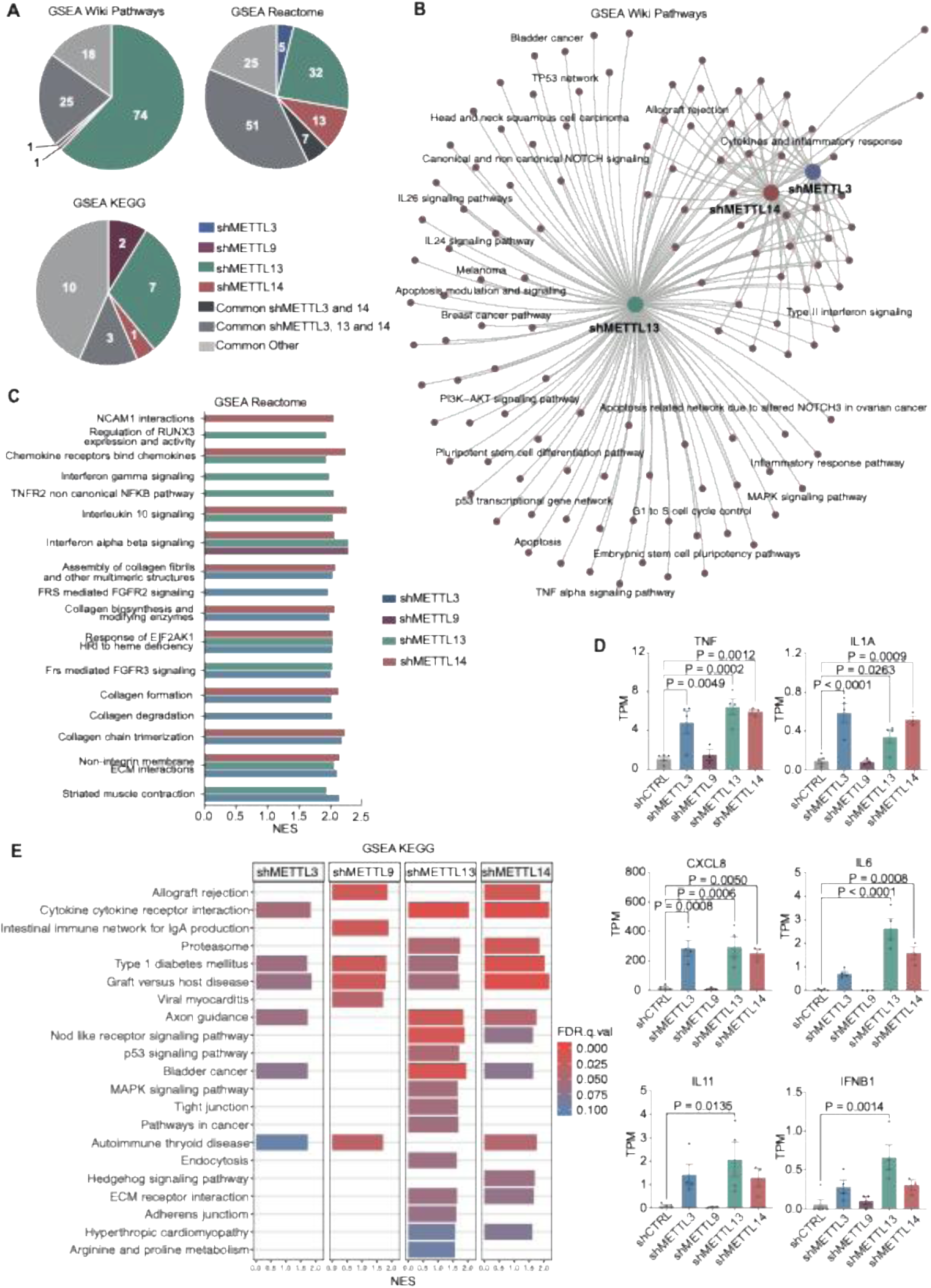
METTL3, METTL13 and METTL14 converged in regulating immune signaling, with many distinct pathway alterations caused only by METTL13. A) Distribution of significantly enriched pathways in each shMETTL compared to shCTRL generated through GSEA. Created in GSEA and MSigDB, FDR q<0.1, including only protein-coding genes, using three different gene set libraries (Wiki Pathways, Reactome and KEGG). Groups are based on which pathways are uniquely altered by shMETTL3 (n=4), shMETTL9 (n=3), shMETTL13 (n=4) or shMETTL14 (n=3), as well pathways altered in several conditions; the m6A writer complex (METTL3 and METTL14) with and without METTL13, as well as all other combined conditions, compared to shCTRL (n=5) in human CD34^+^ HSPCs. B) Network plot of GSEA Wiki Pathways in shMETTL3, shMETTL13 and shMETTL14 compared to shCTRL (FDR q<0.1). C) Top 10 significant GSEA Reactome pathways in shMETTL3, shMETTL9, shMETTL13 and shMETTL14 compared to shCTRL (FDR q<0.1). Results are displayed as normalized enrichment score (NES). D) Dysregulated genes involved in inflammatory signaling (TNF, IL1A, CXCL8, IL6, IL11 and IFNB1) in shMETTL3, shMETTL9, shMETTL13 and shMETTL14 compared to shCTRL. Significance was calculated using ordinary one-way ANOVA with multiple comparisons compared to shCTRL, as well as Dunnett correction, results are displayed as TPM, mean ± SEM. E) GSEA of shMETTL3, shMETTL9, shMETTL13 and shMETTL14 compared to shCTRL using KEGG legacy pathways (FDR q<0.1) Results are displayed as NES and FDR q-value.

METTL13 knockdown exclusively enriched 74 Wiki Pathways, with 25 pathways altered in co-ordinance with METTL3 and METTL14 (Fig. 3B, SF3A). Unique gene sets affected by METTL13 knockdown included several pathways controlling apoptosis and stem cell pluripotency, as well as genes associated with different cancer types, such as melanoma and breast cancer (Fig. 3B, SF3D). Top Reactome gene sets included several immune regulatory pathways, altered by METTL3, METTL9, METTL13 and METTL14 either together or separately (Fig. 3C). METTL13 and METTL14 converged in regulating chemokine receptor binding and IL10 signaling, and together with METTL9 altered IFNα and IFNβ signaling. Analysis of differentially expressed genes revealed alterations to cytokines and chemokines, further endorsing the role of METTL-family members in regulating proper immune signaling. METTL3, METTL13 and METTL14 knockdown all increased levels pro-inflammatory cytokines such as TNF, IL1A and CXCL8 (Fig. 3D) (54–56). METTL13 knockdown increased expression of IL-6 family members IL6 and IL11 as well as IFNβ1a, known to inhibit cell cycle progression in hemopoietic malignances (57). GSEA further revealed dysregulation of several KEGG pathways involved in inflammatory signaling, caused by knockdown of several of the METTL genes, including enrichment of cytokine-cytokine receptor interaction and graft versus host disease (Fig. 3E). Additionally, knockdown of METTL13 had the most exclusively dysregulated KEGG pathways, several crucial for malignant transformation, such as p53 signaling and pathways in cancer.

These results reveal that most altered pathways were either affected by the m6A writer complex together with METTL13 or by METTL13 alone. This suggests that all four METTLs converge in regulating immune signaling in HSPCs. In contrast, the distinctively impacted pathways indicate METTL13 may be uniquely disposed to promote survival, proliferation and possibly oncogenic transformation of HSPCs.

### METTL13 knockdown altered pathways involved in malignant transformation and cancer progression

The unique gene expression program following METTL13 knockdown led us to further delineate the specific role of METTL13 in HSPCs. Inhibition of METTL13 led to altered expression of many genes involved in important cellular processes, including upregulation of CD70, a member of the TNF superfamily that is tightly regulated during hematopoiesis, which together with CD27 plays a role in leukemia stem cell proliferation and survival (Fig. 4A, SF4A) (58). METTL13 knockdown also downregulated CEACAM6, which has been shown to promote tumor proliferation and migration through ERK-MAPK and PI3K-AKT signaling pathways in several cancers, including lung and colon cancer (59).

**Figure 4.**
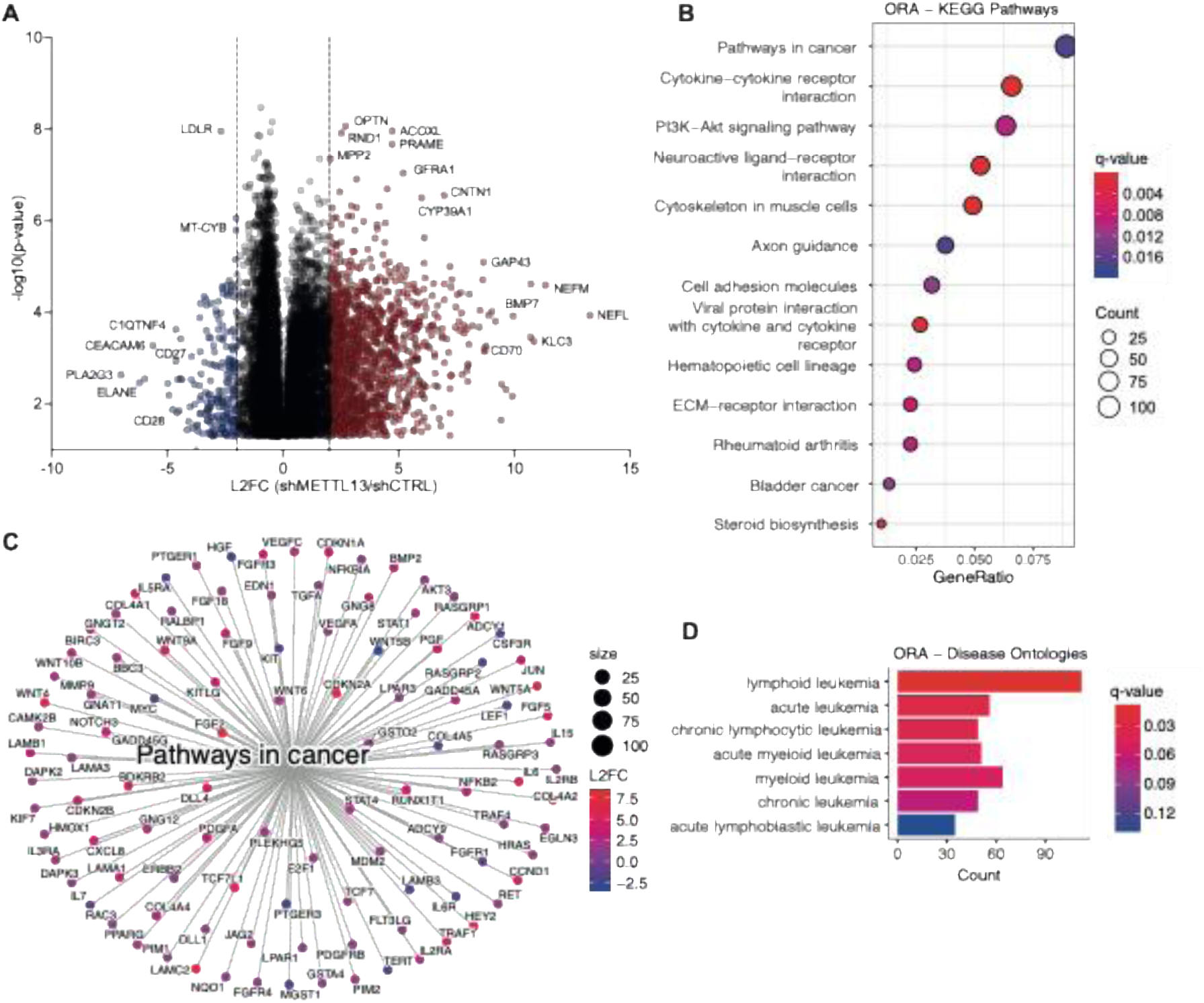
METTL13 knockdown caused dysregulation of pathways involved in malignant transformation and cancer progression. A) Volcano plot of differentially expressed genes following METTL13 knockdown (shMETTL13, n=4) compared to the control (shCTRL, n=5) in human CD34^+^ HSPCs. Significance was calculated using unpaired two-tailed t-test, results are displayed as L2FC and negative log10 p-value (p <0.05). B) ORA of the top enriched KEGG pathways in shMETTL13 compared to shCTRL. Created in R, with packages clusterprofiler and enrichPlot, statistics was set to: q.value <0.1, L2FC cutoff = 1, p-adjust method = BH. C) Network plot of the KEGG gene set pathways in cancer in shMETTL13 compared to shCTRL, nodes are colored by L2FC. D) ORA of disease ontologies focused on hematological malignances. Created in R, packages with clusterprofiler, DOSE and enrichplot, statistics were set to: q.value ≤ 0.1, L2FC cutoff = 1, p-adjust method = BH.

Over-representation analysis (ORA) revealed dysregulation of several KEGG pathways following METTL13 knockdown (Fig. 4B). Many pathways were involved in inflammatory signaling, including TNF signaling and cytokine-cytokine receptor interaction, as well important pathways regulating cell survival, such as PI3K-Akt signaling (Fig. 4B). The pathway with the greatest number of dysregulated genes was pathways in cancer, including many genes crucial for oncogenic progression, which belong to key regulatory processes such as the WNT pathway and NOTCH signaling (Fig. 4B, 4C).

Moreover, ORA of METTL13 knockdown revealed involvement of METTL13 in various disorders, altering numerous disease ontologies, many involving infection and inflammatory reactions, like encephalitis and pneumonia (Fig. SF4B). METTL13 knockdown also altered genes belonging to disease ontologies involved in hematological malignancies, mainly lymphoid leukemias and acute leukemias, as well as several myeloid and chronic leukemias (Fig. 4D). These results further strengthen the hypothesis that METTL13 might have distinct molecular functions regulating malignant alterations in HSPCs, potentially inducing pre-leukemic transformation.

### Upregulation of METTL13 was associated with a high-risk profile and a decreased probability of survival in ALL

Our initial investigation of changes to the genetic profile of HSPCs following knockdown of METTL-family members suggest a potentially unique role of METTL13 in the context of leukemia transformation. To investigate this possibility, we focused on pediatric acute lymphoblastic leukemia (ALL). Indeed, many of the top mutated genes in pediatric T-ALL were downregulated following METTL3, METTL13 and METTL14 knockdown, such as TAL1, LEF1, PHF6, and PTEN (Fig. 5A) (60). Similarly, knockdown caused a loss of ETV6 and CREBBP, candidate driver genes in B-ALL (Fig. SF5A). In contrast, both CDKN2A, a gene with tumour suppressive potential in both T-ALL and B-ALL, as well as KRAS, a B-ALL driver gene, were upregulated (61) (Fig. SF5A). Knockdown of all three METTLs caused a decreased expression of surface marker CD34, mainly found on HSCs, as well as on leukemia-inducing stem cells (Fig. SF5A) (62). Additionally, FBXW7 and FOXO3 were only modified by METTL13 knockdown, both which were upregulated by the loss of METTL13. FBXW7 and FOXO3 have been identified as negative regulators of T-ALL progression, the former inhibiting NOTCH1 activity and the latter inducing apoptosis in pediatric T-ALL (Figure 5B) (63, 64).

**Figure 5.**
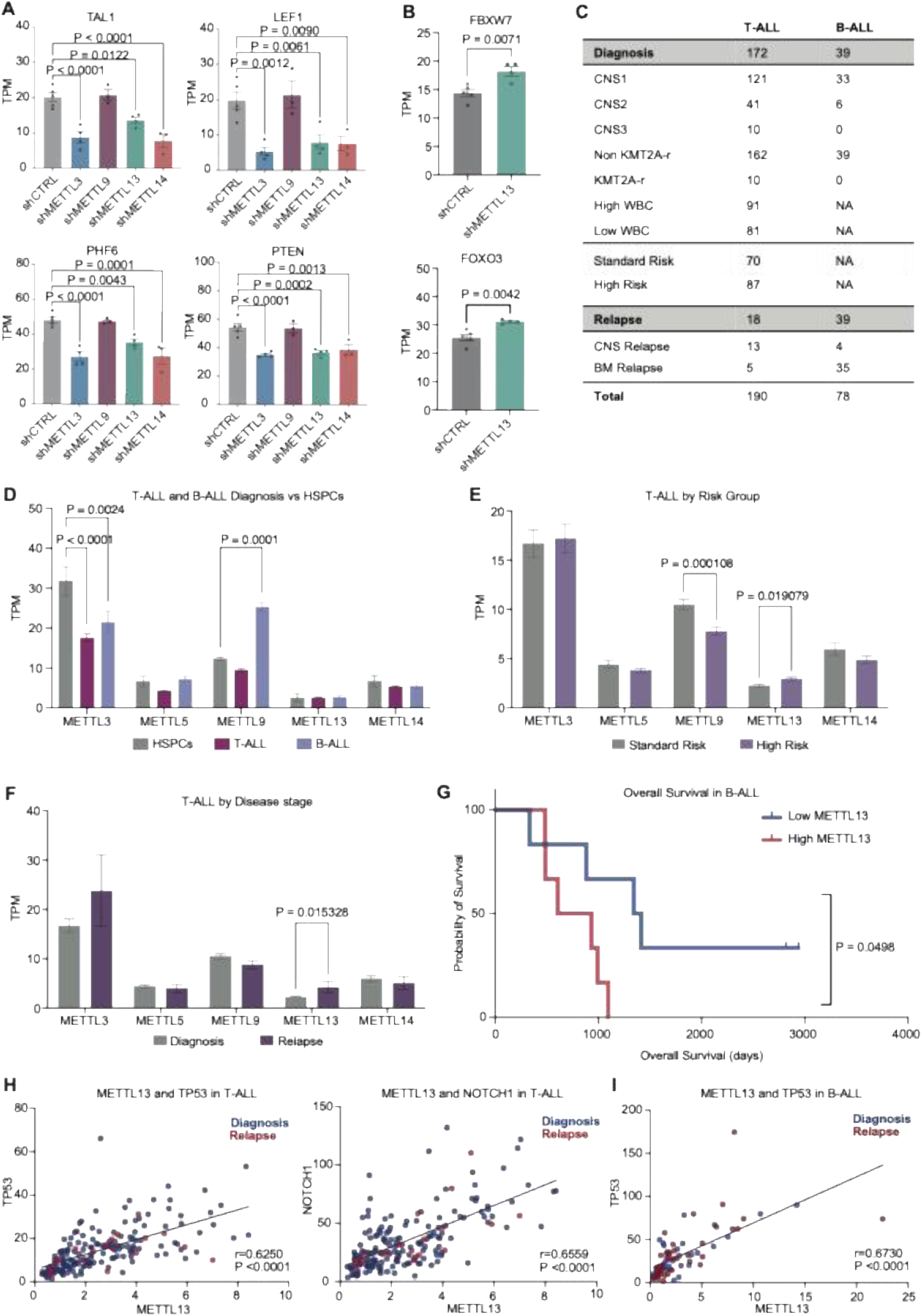
Upregulation of METTL13 associated with a high-risk ALL profile and a decreased probability of survival. A) Dysregulated genes (TAL1, LEF1, PHF6 and PTEN) following METTL3 (n=4), METTL9, (n=3) METTL13 (n=4) and METTL14 (n=3) knockdown in human CD34^+^ HSPCs compared to the control (shCTRL, n=5). Significance was calculated using ordinary one-way ANOVA with multiple comparisons compared to shCTRL, as well as Dunnett correction, results are displayed as TPM, mean ± SEM. B) Uniquely dysregulated genes (FBXW7 and FOXO3) in shMETTL13 (not affected by METTL3, METTL9 or METTL14 knockdown) compared to shCTRL. Significance was calculated using unpaired two-tailed t-test, results are displayed as TPM, mean ± SEM. C) Table of ALL patient characteristics and subgroups used for RNA-sequencing analysis (publicly available by the TARGET Initiative). Samples were divided into T-ALL (n=190) or B-ALL (n=78), diagnosis or relapse, as well as different high-risk factors (CNS infiltration, KMT2A-r and WBC). D) Expression levels of METTL3, METTL5, METTL9, METTL13 and METTL14 in T-ALL (n=162) and B-ALL (n=39) diagnosis samples (publicly available by the TARGET Initiative) compared to normal HSPCs (CD34+ CB, n=5) obtained through RNA-sequencing. Significance was calculated using multiple unpaired t-test, results are displayed as TPM, mean ± SEM. E) Expression levels of METTL3, METTL5, METTL9, METTL13 and METTL14 in T-ALL patient samples (publicly available by the TARGET Initiative) generated through RNA-sequencing, displayed as TPM, mean ± SEM. T-ALL samples were grouped by risk stratification, into standard risk (CNS negative, non KMT2A-r and low WBC, n=70), and high risk (CNS-infiltrated, KMT2A-r or high WBC, n=87). Significance was calculated using multiple unpaired t-test. F) Expression levels of METTL3, METTL5, METTL9, METTL13 and METTL14 in T-ALL patient samples (publicly available by the TARGET Initiative) generated through RNA-sequencing, displayed as TPM, mean ± SEM. T-ALL samples were grouped by disease stage, into diagnosis (standard risk, n=87) and relapse samples (BM relapse, n=5). Significance was calculated using multiple unpaired t-test. G) Survival probability in B-ALL patients samples (publicly available by the TARGET Initiative) by METTL13 expression levels (top and bottom 15%, n=6 in each group). Significance was calculated using Log-rank (Mantel-Cox) test. H) Correlation of METTL13 and TP53 as well as METTL13 and NOTCH1 in T-ALL patient samples, colored by the disease stage (diagnosis = blue, n=162 and relapse = red, n=18), results are displayed as TPM. Correlation was calculated using Pearson correlation coefficients with a two-tailed with 95% confidence interval. I) Correlation of METTL13 and TP53 in B-ALL patient samples, colored by disease stage (diagnosis = blue (n=39), relapse = red (n=39)), results are displayed as TPM. Correlation was calculated using Pearson correlation coefficients with a two-tailed with 95% confidence interval.

To further evaluate the potential involvement of METTL-family members in ALL progression, we performed an RNA-sequencing analysis on pediatric ALL samples, pre-generated by the TARGET initiative (Fig. 5C, Supplemental table 3). For this, a total of 190 T-ALL patients and 78 B-ALL patients were included, with samples obtained both at the time of diagnosis and following relapse. Our analysis identified a significant loss of METTL3 in both T- and B-ALL but did not identify any significant differences in METTL13 expression compared to healthy HSPCs (Fig. 5D). Clinically, ALL diagnosis is based on several risk factors, including white blood cell count (WBC), chromosomal rearrangements such as KMT2A-rearrangement (KMT2A-r) and central nervous system (CNS) infiltration, categorizing patients into different risk groups (65). Among T-ALL samples, we identified an upregulation of METTL13 in both high risk (CNS-infiltrated, KMT2A-r or a WBC above 100) and relapsed samples (with bone-marrow origin) compared to standard risk diagnosis samples (CNS negative, non-KMT2A-r, and a WBC below 100) (Fig. 5E, 5F). This type of risk stratification was not observed for other METTL genes, including METTL3 or METTL14. In B-ALL, although no difference in METTL13 expression was identified between the samples at the time of diagnosis or at relapse, low METTL13 associated with a higher probability of survival (Fig. 5G, SF5B). In line with results obtained after METTL13 knockdown in HSPCs, METTL13 expression correlated positively with TP53 expression in both B- and T-ALL, as well as with NOTCH1 in T-ALL (Fig. 5h, 5I). Our findings indicate converging functions of m6A writer genes METTL3 and METTL14, as well as METTL13, in promoting expression of oncogenic driver genes in both B- and T-ALL. In contrast, METTL13 alone was associated with a high-risk profile in T-ALL and a decreased probability of survival in B-ALL.

### Loss of METTL13 impaired T-ALL proliferation and survival

Our gene profiling suggests a potential link between METTL13 expression and leukemia development. However, the role and prognostic effect of METTL13 in pediatric leukemia is largely unexplored. Here, we further dived into the mechanistic link of METTL13 to leukemia propagation using T-ALL as a model. We first examined the METTL13 expression in normal peripheral blood mononuclear cells (PBMCs) and five T-ALL cell lines using western blot (Fig. 6A, SF6A). METTL13 was highly expressed in all five T-ALL cell lines, with the highest expression in Jurkat, SUP-T1, and CUTTL1. To directly examine the function of METTL13, we knocked down METTL13 in three T-ALL cell lines with various level of METTL13 expression and monitored cell proliferation rate as well as cell viability (Fig. 6B-D, SF6B). Indeed, METTL13 inhibition significantly reduced proliferation and survival starting from day 7 post-lentiviral transduction. We reported a stronger inhibition of cell viability and proliferation in high-METTL13 expressing cell lines (Jurkat and SUP-T1) compared with the relatively low-METTL13-expressing line (MOLT4), suggesting the cellular response may be dependent on cell-intrinsic METTL13 expression level.

**Figure 6.**
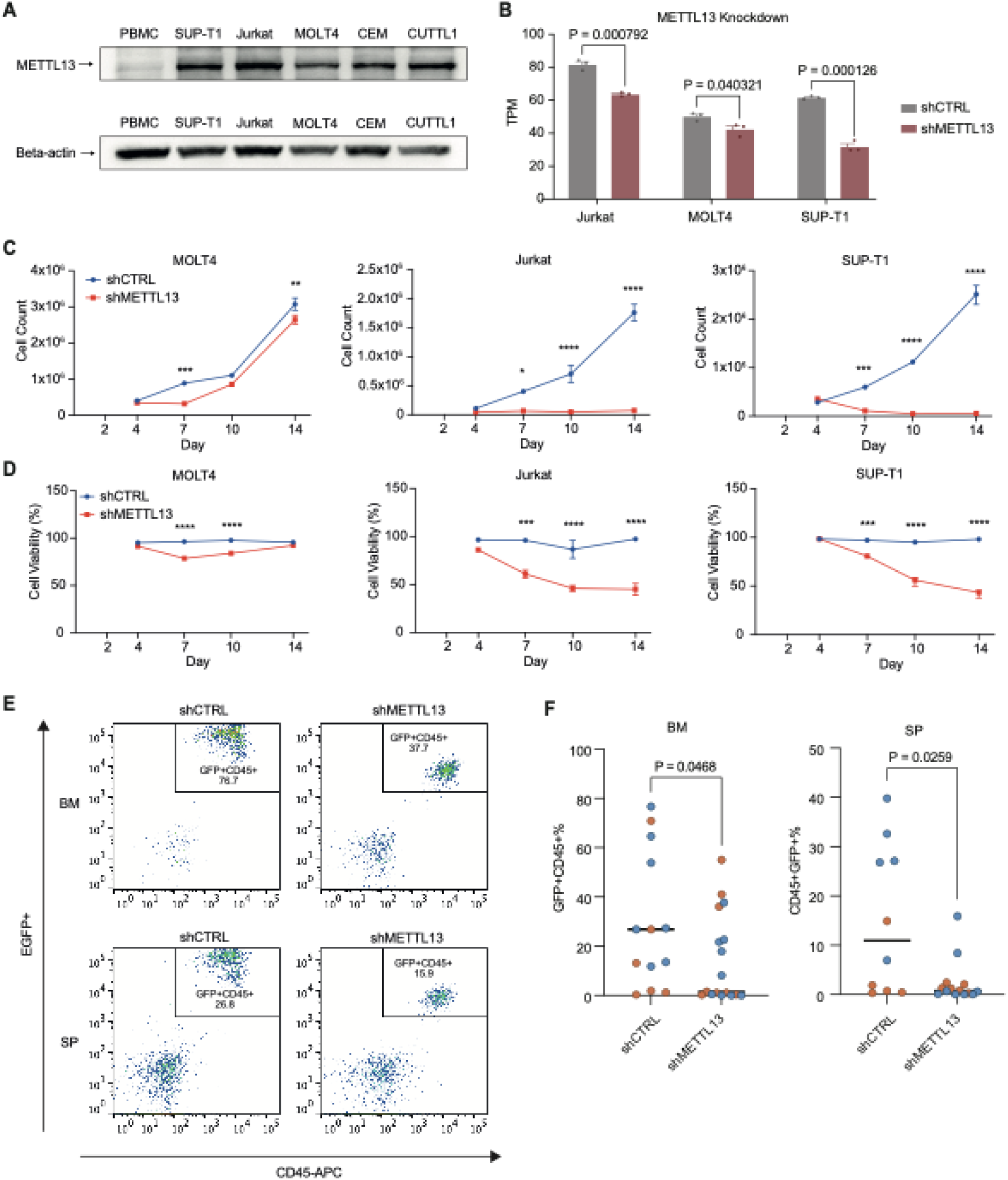
Loss of METTL13 impaired T-ALL cell proliferation and survival. A) Western blot image showing the expression level of METTL13 and beta-actin in normal PBMCs and T-ALL cell lines (SUP-T1, Jurkat, MOLT4, CEM and CUTTL1). B) Validation of METTL13 knockdown (shMETTL13) in T-ALL cell lines Jurkat (n=3), MOLT4 (n=3) and SUP-T1 (n=3) compared to the control (shCTRL, n=3 for each cell line) through RNA-sequencing. Significance was calculated using unpaired two-tailed t-test, results are displayed as TPM, mean ± SEM. C) Total number of viable cells following METTL13 knockdown in T-ALL cell lines MOLT4 (n=3), Jurkat (n=3) and SUP-T1 (n=3) from day 4 to day 14 post transduction compared to the control (n=3 for each cell line). Significance was calculated using two-way ANOVA. D) Cell viability (in percent) following METTL13 knockdown in T-ALL cell lines MOLT4 (n=3), Jurkat (n=3) and SUP-T1 (n=3) from day 4 to day 14 post transduction compared to the control (n=3 for each cell line). Significance was calculated using two-way ANOVA). E) Representative flow cytometry showing human EGFP^+^CD45^+^ leukemia engraftment in NSG-SGM3 mice transplanted with SUP-T1 cells. F) Engraftments of human EGFP^+^CD45^+^ were quantified by flow cytometry in bone marrow (BM) and spleen (SP) of SUP-T1 (blue) and MOLT4 (orange) transplanted mice (n = 10-17 mice per condition).

To validate the *in vitro* results, we also performed an *in vivo* cell line-derived xenograft (CDX) assay. We transduced SUP-T1 and MOLT4 cells with EGFP^+^ scramble control shRNA or a shRNA targeting METTL13 to track the METTL13 knockdown. EGFP+ cells were transplanted into NSG-SGM3 immuno-compromised mice. The leukemia engraftment was detected in both spleen and bone marrow and reached as high as 80% in the control bone marrow (Fig. 6E-F). METTL13 knockdown significantly reduced the leukemia engraftment rate in bone marrow (29.9% in control and 14.4% in shMETTL13) and spleen (15.6% in control and 2.4% in shMETTL13) (Fig. 6F, mean values). Thus, we confirmed that METTL13 is important for leukemia proliferation in both *in vitro* and *in vivo* T-ALL models.

### Knockdown of METTL13 promoted pathways that induce apoptosis and suppress DNA synthesis in T-ALL cells

In addition to the functional validation identifying METTL13 as a regulator of T-ALL proliferation, we wanted to study aspects of METTL13 on a transcriptional level in T-ALL. Thus, we performed RNA-sequencing on MOLT-4, Jurkat, and SUP-T1 cell lines following METTL13 knockdown. Knockdown of METTL13 caused alterations to thousands of genes within each cell line, both up- and downregulated (Fig. SF7A, Supplemental table 4). The three T-ALL cell lines clustered separately in PCA with most differentially expressed genes unique to each cell line, indicating a greater variability between the cell lines, which likely represents the heterogenicity of T-ALL (Fig. SF7B, SF7C). Despite this, all three cell lines also clustered into distinct METTL13 knockdown and control groups. The three cell lines shared only 1,529 significantly differentially expressed genes following METTL13 knockdown (Fig. SF7C-E), highlighting the importance of intercellular variability and cell-specific m6A modifications.

To further study effects of METTL13 knockdown in T-ALL and to counteract the cell-intrinsic differences among cell types, the cell lines were treated as biological replicates, divided into shMETTL13 and shCTRL. PCA clearly showed a separation between the two conditions, which was further reinforced by visualization of the gene expression pattern (Fig. 7A-B). Our analysis revealed that approximately 1% of all genes (a total of 711 genes) were differentially expressed, with the majority being downregulated (Fig. 7C-D). Among affected genes, we identified an upregulation of MDM2, a similar effect recognized in HSPCs with reduced METTL13 levels (Fig. 7E-F). Loss of METTL13 caused downregulation of NRAS, a B-ALL candidate driver gene, and ERBB3, which is frequently overexpressed in cancer, involved in promoting tumor progression (60, 67). Moreover, METTL13 knockdown induced expression of BTG1, identified to induce cell cycle arrest and inhibit cell proliferation in several different malignancies (68).

**Figure 7.**
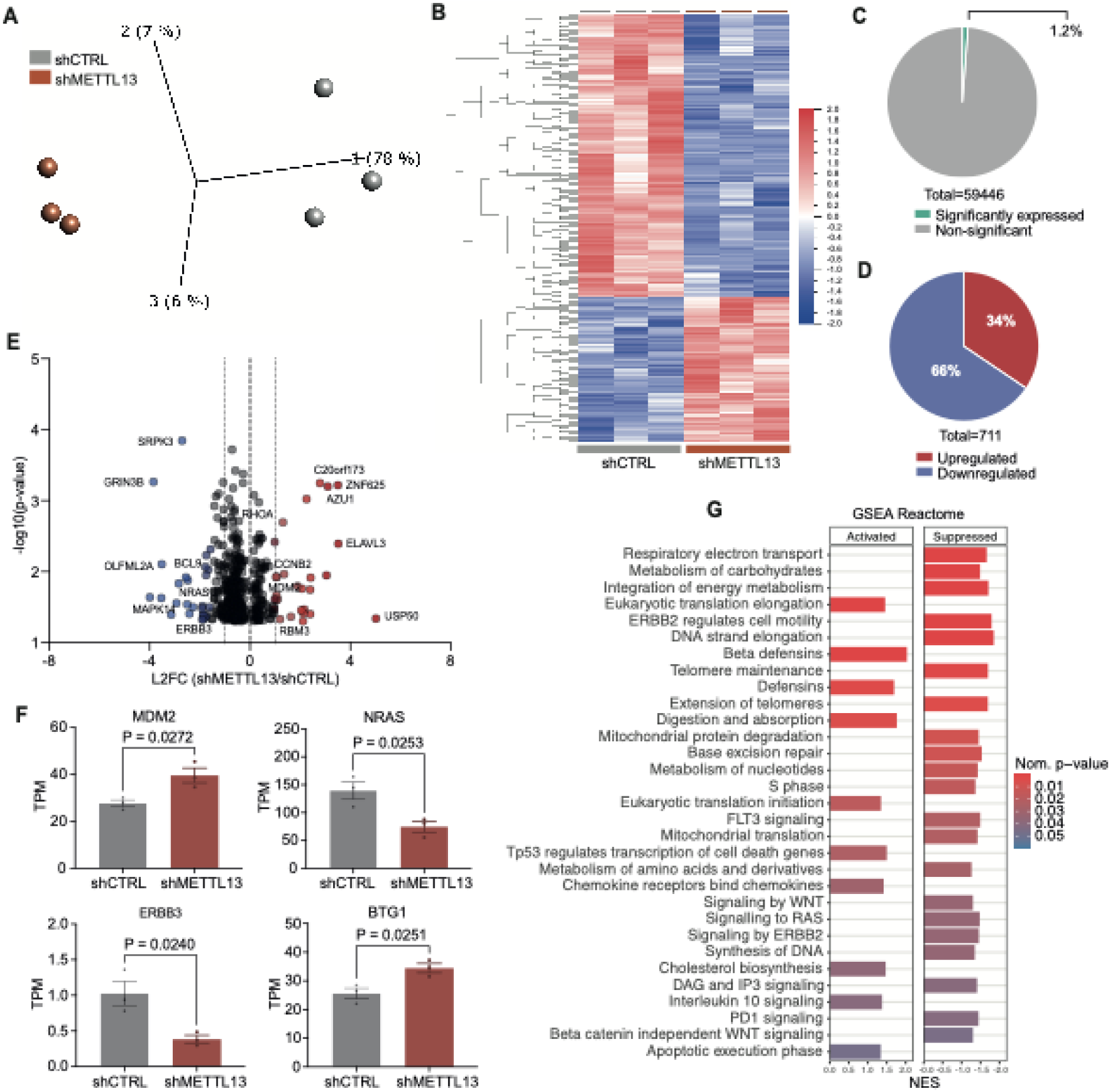
Knockdown of METTL13 promoted pathways that induce apoptosis and suppress DNA synthesis in T-ALL cells. A) PCA plot of T-ALL cell lines as biological replicates following METTL13 knockdown (shMETTL13, n= 3) compared to the control (shCTRL, n=3). Significance was calculated using unpaired two-tailed t-test, p<0.05. B) Heat map of the top 400 differentially expressed genes looking at T-ALL cell lines as biological replicates in shMETTL13 (n=3) compared to the shCTRL (n=3). Significance was calculated using unpaired two-tailed t-test, p<0.05. C) Pie chart of the percentage of significantly expressed genes (p<0.05) in T-ALL cell lines as biological replicates (n=3) in shMETTL13 (n=3) compared to shCTRL (n=3). Significance was calculated using unpaired two-tailed t-test, p<0.05. D) Pie chart of the distribution of upregulated versus downregulated significantly expressed genes in T-ALL cell lines as biological replicates in shMETTL13 (n=3) compared to the shCTRL (n=3). Significance was calculated using unpaired two-tailed t-test, p<0.05. E) Volcano plot of significantly expressed genes (p<0.05) in T-ALL cell lines as biological replicates in shMETTL13 (n=3) compared to the shCTRL (n=3). Only protein-coding genes were included in this plot. Significance was calculated using unpaired two-tailed t-test, results are displayed as L2FC and negative log10 p-value (p <0.05). F) Dysregulated genes (MDM2, NRAS, ERBB3 and BTG1) in shMETTL13 (n=3) compared to shCTRL (n=3 Significance was calculated using unpaired two-tailed t-test, p<0.05, results are displayed as TPM, mean ± SEM. G) GSEA of some of the top significantly enriched KEGG pathways (Created in GSEA and MSigDB, nominal p-value <0.05) in shMETTL13 (n=3) compared to shCTRL (n=3) in T-ALL cells, using only protein-coding genes, displayed as activated (positive NES) or suppressed (negative NES).

Finally, GSEA of Reactome pathways revealed that METTL13 knockdown led to changes in several signaling pathways and metabolic regulation (Fig. 7G). Loss of METTL13 caused suppressed nucleotide metabolism, WNT and RAS signaling as well as DNA synthesis. Contrarily, METTL13 knockdown activated TP53 regulation of cell death genes, chemokine receptor binding to chemokines, IL10 signaling and apoptotic execution phase. The results from the pathway analysis, together with both the *in vitro* and *in vivo* validation, highlight the potential impact METTL13 could have on leukemia cell survival and proliferation. Collectively, these data indicate a possible oncogenic role of METTL13 in pediatric T-ALL pathogenesis.

## Discussion

To better understand the role of m6A methylation and A-to-I editing, including their distinct effects and potential crosstalk, we investigated these RNA modifications in the context of normal HSCs as well as in malignant transformation leading to pediatric leukemia. We have shown that ADAR1 has an overall inhibitory effect on the m6A complex in HSPCs, yet promotes the expression of METTL13, whose role in oncogenic transformation remains undefined. Our results uncover a novel role for METTL13 in regulating gene expression in HSPCs, both in co-ordinance with the m6A writer complex, and with unique effects on genes involved in apoptosis and cancer progression. We have also identified that METTL13 regulates T-ALL cell survival and proliferation both *in vitro* and *in vivo* and was associated with a high-risk profile in pediatric T-ALL patient samples.

Previous studies have shown that RNA methylation by the m6A complex induced RNA editing by increasing ADAR1 levels in response to IFN stimulation (42) and METTL3 was found to increase protein levels of ADAR1 in glioblastoma (43). In contrast, ADAR1 was found to preferentially bind to m6A-depleted RNA transcripts, indicating a negative correlation between A-to-I editing and m6A, where m6A suppresses A-to-I editing (69). In this study, we revealed a general suppressive effect of ADAR1 on members of the m6A complex on a transcriptional level in human HSPCs. ADAR1 overexpression inhibited expression of the main writer genes METTL3 and METTL14, as well as both known erasers FTO and ALKBH5. In contrast, an orphan METTL-family member, METTL13, was augmented by the induction of ADAR1. Whether METTL13 acts in co-ordinance with other METTL-family members, or has similar effects on HSC maintenance as ADAR1, will need to be carefully investigated by future studies.

Research on the m6A complex is extensive, and aberrant activity of m6A modifications have been shown to promote development of both solid tumors and hematological malignancies (37). Despite this, the distinct role of each METTL-family member remains undefined. Therefore, we decided to knockdown m6A writers METTL3 and METTL14, along with less studied METTL5, METTL9 and METTL13 in normal HSPCs, to further study this complex epitranscriptomic landscape. The genetic profile generated by METTL knockdown revealed many similarities between METTL3 and METTL14, and surprisingly, METTL13. Most differentially expressed genes were altered by the m6A writer complex as well as by METTL13, whose connection to the m6A complex remains unclear. Knockdown of METTL3, METTL13 and METTL14 modified many genes crucial for malignant development, such as TP53, MDM2 and c-MYC. Moreover, GSEA revealed dysregulation of several pathways following knockdown of METTL3 and METTL14 in HSPCs, many involved in inflammatory signaling. Interestingly, most pathways enriched by METTL3 and METTL14 were also enriched by METTL13 knockdown, further strengthening the notion of METTL13 acting in convergence with the m6A writer complex. Despite many commonly altered pathways, GSEA discovered numerous pathways uniquely altered by METTL13 knockdown, including p53 signaling and several cancer-associated pathways.

Current research has started to shift focus towards METTL13, focusing on understanding its biological function and possible oncogenic potential. Recent reports have identified METTL13 as a lysine methyltransferase that modifies the eEF1a protein at lysine 55, thus promoting translation (70, 71). METTL13 has been recognized to correlate with a decreased survival probability in malignancies such as lung and pancreatic cancer and was found to augment metastasis in gastric cancer (71, 72). Existing studies focus mainly on methylation of eEF1A and changes in translational output, several highlighting the potential oncogenic role of METTL13 in cancer. In contrast, loss of METTL13 was considered a poor prognostic marker in clear cell renal cell carcinoma, inhibiting proliferation and metastatic capacity of cancer cells (73). However, research on METTL13 and its role in HSC biology is sparse. The extensively overlapping gene expression programs between METTL3-METTL14 complex and METTL13 identified in this study indicate that functions of METTL13 align with the m6A pathway in the cellular context of HSCs. Future studies are necessary to systematically de-couple the roles of METTL13 in m6A regulation and protein translation control.

Since METTL13 was recently reported as an eEF1a methyltransferase, it is surprising that the gene expression pattern of METTL13 substantially overlaps with that of m6A writer complex. These results, combined with our data highlighting the effects of METTL13 in malignant transformation, prompted further analysis to unravel its specific role in hematological diseases. Our findings suggest transcriptional changes following knockdown of METTL13 in HSPCs, capable of promoting oncogenic progression, where the main altered gene set following ORA was pathways in cancer. Moreover, analysis of disease ontologies revealed enrichment of many leukemia-related malignancies of both myeloid and lymphoid origins. To date, the role of METTL13 in hematological cancers such as pediatric ALL remains largely unexplored. Therefore, we examined transcriptional alterations to candidate driver genes in pediatric ALL following knockdown of members in the METTL-family in HSPCs. We identified a loss of numerous oncogenic drivers of both B- and T-ALL following knockdown of METTL3, METTL13 and METTL14, including TAL1, LEF1 and CREBBP. Additionally, the expression of inhibitory tumour suppressors, such as CDKN2A, was upregulated.

To further explore the role of METTL-genes in pediatric ALL, we utilized a publicly available RNA-sequencing dataset (generated by the TARGET initiative) and report that METTL13 was overexpressed in both high-risk and relapsed T-ALL. Of note, METTL13 was the only METTL-family member augmented by ADAR1 overexpression in HSPCs, which was recently described to drive relapse in pediatric T-ALL (10).

Finally, we report reduced survival and proliferation of T-ALL cells following METTL13 knockdown, both *in vitro* and *in vivo*. T-ALL cell lines with a higher endogenous METTL13 expression showed a stronger response to METTL13 inhibition. Knockdown of METTL13 also significantly reduced the leukemia engraftment rate in our CDX model. RNA-sequencing analysis and GSEA of T-ALL cells with depleted METTL13 expression revealed activation of both p53 transcriptional regulation of cell death genes and apoptosis. Loss of METTL13 also caused reduced DNA synthesis and inhibition of signaling pathways crucial for cell proliferation and leukemic progression, such as WNT and RAS signaling.

In conclusion, the collective findings following METTL13 knockdown in HSPCs and in T-ALL cells suggest that METTL13 may be involved in promoting pre-leukemic development of HSPCs and in the pathogenesis of T-ALL. Future studies are necessary to functionally determine effects of METTL13 in leukemic progression, yet our findings highlight a novel, distinct role of METTL13 in T-ALL development.

## Materials and Methods

### Cord blood and cell line handling

CD34^+^ human cord blood samples and PBMCs were purchased from commercial vendors (StemCell Technology or AllCells) and stored in liquid nitrogen until ready for use. T-ALL cell lines (CEM, CVCL_0207; CUTTL1, CVCL_4966; MOLT-4, RRID:CVCL_A1BB; SUP-T1, RRID:CVCL_1714; and Jurkat, RRID:CVCL_0065) were purchased from ATCC and maintained in RMPI media (Gibco, 11875119) supplemented with 10% FBS (Gibco) and 1% penicillin-streptomycin (Gibco, 15140) at 37° C in 5% CO2.

### Lentiviral construct and transduction

To determine the function of members of the METTL-family, lentiviral knockdown in HSPCs and T-ALL cells was performed. Lentiviral construct of shRNAs targeting METTLs was purchased from VectorBuilder (Vector ID: VB240522-1552yyd). All lentivirus was tested by transducing HEK293T cells, and the knockdown efficiency and titers were assessed by FACS analysis of GFP signal and RT-qPCR of the target genes. Cord blood CD34^+^ cells were cultured in 96-well plate (5 x10^5^ cells per well) containing StemPro (Life Technologies) media supplemented with human cytokines (IL-6, stem cell factor (SCF), Thrombopoietin (Tpo) and FLT-3, all from R&D Systems) for 2-3 days at lentivirus at a MOI of 100-200. T-ALL cell lines were cultured in 24-well plate or 6-well plate (5-10 x10^5^ cells per well) in culture media with lentivirus at a MOI of 5-10. Cells were then collected for downstream analysis.

### Animal experiments and Flow Cytometry analysis

All mouse studies were conducted under protocols approved by the Institutional Animal Care and Use Committee (IACUC) of the University of California, San Diego and were in compliance with federal regulations regarding the care and use of laboratory animals: Public Law 99-158, the Health Research Extension Act, and Public Law 99-198, the Animal Welfare Act which is regulated by USDA, APHIS, CFR, Title 9, Parts 1, 2, and 3. Immunocompromised NSG-SGM3 mice were bred and maintained in the Sanford Consortium for Regenerative Medicine vivarium according to IACUC approved protocols of the University of California, San Diego. Neonatal mice of both sexes were used in the study. MOLT4 or SUP-T1 cells were injected intrahepatically into 2-3 days old neonatal NSG-SGM3 mice at 2.5-5×10^4^ cells per pup. Leukemic engraftment was quantified by FACS analysis-based peripheral blood screening of human CD45^+^ population starting from week 3 for every week until the engraftment reaches 10%. Mice were humanely sacrificed at 4-8 weeks, and cells were collected from hematological organs (bone marrow and spleen) for FACS analysis. Cells were resuspended in staining media (ice cold DPBS with 2% FBS), followed by blocking using FcR block (Cat 130-059-901, Miltenyi Biotech) for 15 minutes. CD45-APC cell surface antibody (Cat # 304037, Biolegend, San Diego, CA) was added a final dilution of 1:25 and incubated on ice for 30 min in the dark. DAPI solution was added before analysis to exclude dead cell debris. Flow analysis was performed on BD Aria Fusion, Aria II.

### Western Blot analysis

Cells were collected and washed three times with cold PBS and resuspended in RIPA buffer supplemented with protease and phosphatase inhibitors and quantified by BCA assay. Protein lysate (10 μg) was separated by 10% SDS–PAGE and transferred to PVDF membranes. The membrane was blocked in 5% BSA/20mM Tris-HCL for 30 min while rocking and probed with primary antibodies (METTL13 antibody, abcam, ab186002; beta-actin antibody, Millipore Sigma, A2228) overnight at 4 °C followed by incubation with HRP-conjugated secondary antibodies for 2 hours at room temperature. Signals were visualized using SuperSignal West Femto Substract (ThermoFish, #34096) on a ChemiDoc System (Bio-rad).

### RNA extraction and Quantitative Real-Time Polymerase Chain Reaction (RT-qPCR)

For validation of lentiviral knockdown, RNA was extracted and analyzed through RT-qPCR. RNA extraction was performed using RNeasy micro extraction kits (QIAGEN) following the manufacturer’s protocol. A DNase incubation step was included to digest any genomic DNA. RNA (100-1,000 ng) was converted to cDNA using the Super-Script III kit (ThermoFisher Scientific) according to the manufacturer’s recommended protocol. qRT-PCR was performed using SYBR GreenER Super Mix (Life Technologies) on BioRad CFX382 with 5 ng of template mRNA and 0.2 μM of each forward and reverse primer. Target mRNA was normalized to hypoxanthine phosphoribosyl transferase (HPRT) mRNA transcript levels and fold change was calculated via the delta-delta cycle threshold (CT) method. RT-qPCR primers are listed in supplemental table 5.

### Viability and proliferation analysis

To define effects of METTL13 knockdown on T-ALL cells, T-ALL cell lines were transduced with lentiviral scramble control or shRNA targeting shMETTL13 at a low MOI of 10-20 in culture media without pen/strep. After 2-3 days, cells were plated in 96-well plates at 1 x10^5^ cells per well. At each time point, cell number and viability were determined by trypan blue assay.

### RNA-sequencing following lentiviral knockdown

The RNA-sequencing dataset transduced with ADAR1 overexpression in HSPCs was available from previous studies (BioProject: PRJNA319866) (6). For RNA-sequencing of METTL knockdown in HSPCs and T-ALL, samples with RNA integrity numbers (RIN) ≥7 were proceeded for bulk RNA-sequencing. RNA-sequencing of HSPCs transduced with METTL3, METTL5, METTL9, METTL13 and METTL14 knockdown was performed using 50-100 ng of RNA by Scripps Research on SMARTer seq system, with paired-end 150 reads, at 50 million reads per sample. Obtained reads were aligned using STAR two-pass alignment method and the reference genome GRCh38.84 together with the corresponding GFT file, to generate transcriptome-coordinate based BAM files as described in a previous study (10). Raw counts were obtained through STAR, using the ENCODE STAR-RSEM pipeline generating the numbers of reads aligned to each gene. Transcripts per million (TPM) values were calculated over the total collapsed exonic regions for each gene. RNA-sequencing of T-ALL cells transduced with METTL13 knockdown was performed using 100 ng of RNA by Novogene on NovaSeq X Plus sequencing system with paired-end 150 reads. Raw counts were generated from BAM files together with the provided GTF file in R (version 4.3.1) using FeatureCount from the Rsubread package (version 2.16.1), with the pipeline set to unstranded and paired end reads. Raw count data was normalized in R (version 4.3.1) to TPM using the mRNA expression transformation guideline provided by the GDC bioinformatics pipeline.

### Publicly available RNA-sequencing downloading and normalization

RNA-sequencing data from pediatric T-ALL patients was obtained from the publicly available data generated by the TARGET initiative. The RNA-sequencing dataset (.BAM) were acquired via the GDC Data Portal (https://portal.gdc.cancer.gov/projects/TARGET-ALL-P2). A total of 190 bone marrow-derived T-ALL samples (192 from time of diagnosis, 18 after relapse, non-longitudinal) and 78 bone marrow-derived B-ALL samples (39 from time of diagnosis, 39 after relapse, longitudinal) were included. Sample inclusion was based on the sample type (bone marrow), disease status (diagnosis and relapse) as well as MLL status (positive or negative, samples with status unknown were excluded). Raw counts were generated using the SeqMonk Mapped Sequence Data Analyzer tool (version 1.47.1). The RNA-seq quantitation pipeline was set to follow the subsequent criteria: transcript features were set to mRNA, library type was selected as opposing strand specific and transcript isoforms were merged. Raw count data from CD34^+^ cord blood was obtained from the publicly available data set found in the Gene Expression Omnibus (GEO) at GSE190269 (74). Raw count data was normalized in R (version 4.3.1) to TPM using the mRNA expression transformation guideline provided by the GDC bioinformatics pipeline.

### RNA quantification and functional enrichment

Subsequent analysis was based on normalized RNA-sequencing data (TPM), generated from raw counts as described above. PCA and heatmaps were created using Qlucore Omics Explorer (version 3.9). Significance for the heatmap was calculated using multi-group ANOVA (q<0.1, SD<0.05). ORA was performed by including all significantly differentially expressed genes (calculated using unpaired two-tailed t-test, p < 0.05) and generated in R (version 4.3.1), using packages ClusterProfiler (version 4.8.3), DOSE (version 3.28.2) and Enrichplot (version 1.20.3). For ORA, p-adjust method was set to BH and L2FC cutoff of 1. GSEA was performed using GSEA software (version 4.3.2) provided by Broad Institute and UC San Diego (75, 76), utilizing human KEGG Legacy (version 2024.1.Hs), Reactome (version 2024.1.Hs) and Wiki Pathway (version 2024.1.Hs) gene sets. For GSEA, only protein-coding genes were included. For analysis of differentially expressed genes, genes with a L2FC = NA were excluded. Differentially expressed genes were obtained using unpaired two-tailed t-test between the different experimental conditions compared to the control group (p < 0.05).

### Statistical analysis

Statistical analyses were performed using ordinary one-way ANOVA of the mean of every condition against the control group with multiple comparisons (using normalized TPM values), with Dunnett correction or using unpaired two-tailed t-test when comparing normalized TPM values between two groups. Correlation was determined by Pearson correlation coefficients, two-tailed with 95% confidence interval. Analysis of cell viability and proliferation following *in vitro* analysis was performed using ordinary two-way ANOVA with multiple comparisons and analysis of cell engraftment *in vivo* was performed using Mann-Whitney test. The statistical analyses were performed using GraphPad Prism (v9) or R (version 4.3.1). Results are presented as the mean ± SEM. Significance was set to P < 0.05.

## Data availability

The RNA-sequencing dataset used in this study will be uploaded to an appropriate data portal upon request with a special password for editors and reviewers. The data will be made publicly available upon acceptance for publication. Further information and requests for resources and reagents should be directed to and will be fulfilled by Dr. Frida Holm.

## Author Contributions

S.E., A.R.A., I.S., C.L., C.T., Q.J., and F.H. collected and analyzed the data. Q.J., F.H. and S.E. conceptualized the project, analyzed data, and wrote the paper. S.S.F., A.N., and O.H. contributed with scientific expertise and were involved in writing and reviewing the paper.

## Acknowledgments

This work was supported by the Swedish Childhood Cancer Foundation/Barncancerfonde (F.H: TJ2014-0014, PR2017-0086, O.H: PR2021-0112-2), Åke Wiberg Foundation (M18-0151) and Märta och Gunnar V. Philipson Foundation (A.N, F.H), Leukemia Research Foundation (Q.J.), Hartwell Foundation (Q.J.), American Society of Hematology (Q.J.), CureBound Discovery Grant (Q.J.) NIH NCI 1R01CA282792-01A1 (Q.J. ) and NIH NCI 1R03CA287274-01 (Q.J.).

The TARGET Initiative (RNA-sequencing dataset): The results published here are in whole or part based upon data generated by the Therapeutically Applicable Research to Generate Effective Treatments (TARGET) initiative, phs000218, managed by the NCI. The data used for this analysis are available at https://portal.gdc.cancer.gov/projects/TARGET-ALL-P2. Information about TARGET can be found at http://ocg.cancer.gov/programs/target. Part of the data handling was enabled by resources at project number SNIC-2021/22-48 provided by the Swedish National Infrastructure for Computing (SNIC) partially funded by the Swedish Research Council through grant agreement no. 2018-05973.

## Supplemental Figures

**Supplemental Figure 1.**

A) Validation of ADAR1 overexpression in human CD34^+^ HSPCs (n=3) compared to the backbone control (pCDH, n=3). Significance was calculated using unpaired two-tailed t-test, results are displayed as TPM, mean ± SEM.

B) Expression levels of genes in the METTL-family following ADAR1 overexpression compared to the control. Significance was calculated using unpaired two-tailed t-test, results are displayed as TPM, mean ± SEM.

**Supplemental Figure 2.**

A) Validation of METTL knockdown (shMETTL3 (n=3), sMETTL5 (n=3), shMETTL9 (n=3), shMETTL13 (n=3) and shMETTL14 (n=3)) in human CD34^+^ HSPCs compared to the control (shCTRL, n=3) through RT-qPCR. Significance was calculated using unpaired two-tailed t-test, results are displayed as expression level relative to the housekeeping gene (HPRT), mean ± SEM.

B) Validation of METTL knockdown (shMETTL3 (n=4), sMETTL5 (n=2), shMETTL9 (n=3), shMETTL13 (n=4) and shMETTL14 (n=3)) in HSPCs compared to the control (shCTRL, n=5) through RNA-sequencing. Significance was calculated using unpaired two-tailed t-test, results are displayed as TPM, mean ± SEM.

C) PCA plot of shMETTL3, shMETTL5, shMETTL9, shMETTL13 and shMETTL14 compared to shCTRL with the outliers labeled (excluded in sequential analysis). Created in Qlucore Omics Explorer.

**Supplemental Figure 3.**

A) Venn diagram of GSEA Wiki Pathways between shMETTL3, shMETTL9, shMETTL13 and shMETTL14 in human CD34^+^ HSPCs compared to shCTRL (FDR q<0.1).

B) Venn diagram of GSEA Reactome pathways between shMETTL3 (n=4), shMETTL9 (n=3), shMETTL13 (n=4) and shMETTL14 (n=3) compared to shCTRL (n=5) (FDR q<0.1).

C) Venn diagram of GSEA KEGG pathways between shMETTL3, shMETTL9, shMETTL13 and shMETTL14 compared to shCTRL (FDR q<0.1).

D) Bar plot of all unique GSEA Wiki Pathways (FDR.q <0.1) in shMETTL13 compared to shCTRL. Results are displayed by NES.

**Supplemental Figure 4.**

A) Top differentially expressed genes following METTL13 (shMETTL13, n=4) knockdown in human CD34^+^ HSPCs compared to control (shCTRL, n=5). Significance was calculated using unpaired two-tailed t-test (p < 0.05), results are displayed as L2FC (shMETTL13/shCTRL).

B) ORA of top disease ontologies in shMETTL13 compared to shCTRL. Created in R, packages ClusterProfiler, DOSE and Enrichplot, with statistics set to: q.value ≤ 0.1, L2FC cutoff = 1, p-adjust method = BH.

**Supplemental Figure 5.**

A) Dysregulated genes (ETV6, CREBBP, KRAS, CD34 and CDKN2A) METTL3 (shMETTL3, n=4), METTL9, (shMETTL9, n=3) METTL13 (shMETTL13, n=4) and METTL14 (shMETTL14, n=3) knockdown in human CD34^+^ HSPCs compared to the control (shCTRL, n=5). Significance was calculated by ordinary one-way ANOVA with multiple comparisons compared to shCTRL, as well as Dunnett correction, results are displayed as TPM, mean ± SEM.

B) Expression levels of METTL3, METTL5, METTL9, METTL13 and METTL14 in B-ALL patient samples (publicly available by the TARGET Initiative) generated through RNA-sequencing. B-ALL samples were grouped by disease stage, into diagnosis (n=39) or relapse samples (n=39), samples are longitudinal. Significance was calculated using multiple unpaired t-test, results are displayed as TPM, mean ± SEM.

**Supplemental Figure 6.**

A) Western blot image showing the expression level of METTL13 and beta-actin in normal PBMCs and T-ALL cell lines (SUP-T1, Jurkat, MOLT4, CEM and CUTTL1).

B) Validation of METTL13 knockdown (shMETTL13) in T-ALL cell lines Jurkat (n=1), MOLT4 (n=1) and SUP-T1 (n=1) compared to the control (shCTRL, n=1 for each cell line) through RT-qPCR. Significance was calculated using unpaired two-tailed t-test, results are displayed as expression level relative to the housekeeping gene (HPRT), mean ± SEM.

**Supplemental Figure 7.**

A) Distribution of differentially expressed genes in shMETTL13 compared to shCTRL in each T-ALL cell line separately (n=3 for each cell line and condition). Significance was calculated using unpaired two-tailed t-test, p < 0.05.

B) PCA plot of shMETTL13 compared to shCTRL in T-ALL cell lines Jurkat, MOLT4 and SUP-T1 (n=3 for each cell line and condition). Created in Qlucore Omics Explorer.

C) Venn diagram of all differentially expressed genes (p<0.05) in shMETTL13 compared to shCTRL in each T-ALL cell line separately (n=3 for each cell line and condition). Significance was calculated using unpaired two-tailed t-test, p < 0.05.

D) Venn diagram of significantly upregulated genes (p<0.05) in shMETTL13 compared to shCTRL in each T-ALL cell line separately (n=3 for each cell line and condition). Significance was calculated using unpaired two-tailed t-test, p < 0.05.

E) Venn diagram of significantly downregulated genes (p<0.05) in shMETTL13 compared to shCTRL in each T-ALL cell line separately (n=3 for each cell line and condition). Significance was calculated using unpaired two-tailed t-test, p < 0.05.

## Supplemental Tables

**Supplemental Table 1.** Differentially expressed genes following ADAR1 overexpression in HSPCs

**Supplemental Table 2.** Differentially expressed genes following METTL3, METTL5, METTL9, METTL13 and METTL14 knockdown in HSPCs

**Supplemental Table 3.** Clinical characteristics of ALL patient samples provided by TARGET

**Supplemental Table 4.** Differentially expressed genes following METTL13 knockdown in T-ALL cell lines

**Supplemental Table 5.** RT-qPCR Primers

